# A spatially regulated GTPase cycle of Rheb controls growth factor signaling to mTORC1

**DOI:** 10.1101/472241

**Authors:** Marija Kovacevic, Christian H. Klein, Lisaweta Roßmannek, Antonios D. Konitsiotis, Angel Stanoev, Astrid U. Kraemer, Philippe I. H. Bastiaens

## Abstract

Growth factors initiate anabolism by activating mechanistic target of rapamycin complex 1 (mTORC1) via the small GTPase Rheb. We show that the GTPase cycle of Rheb is spatially regulated by the interaction with its GDI-like solubilizing factor (GSF) – PDEδ. Arl2-GTP mediated localized release of cytosolic Rheb-GTP from PDEδ deposits it onto perinuclear membranes where it forms a complex with mTORC1. The membrane associated GTPase activating protein (GAP) TSC2 hydrolyzes Rheb-GTP, weakening the interaction with mTOR. Rheb-GDP is readily released into the cytosol where it is maintained soluble by interaction with PDEδ. This solubilized Rheb is re-activated by nucleotide exchange to be re-deposited by Arl2-mediated release onto perinuclear membranes. This spatial GTPase cycle thereby enables mTORC1 activation to be solely controlled by growth factor induced inactivation of TSC2. The coupling between mTOR activation and spatially regulated Rheb nucleotide exchange makes growth factor induced proliferation critically dependent on PDEδ expression.

## INTRODUCTION

mTORC1 is a central signaling node that interlinks growth factor signals and the processes that drive cellular growth. This is achieved by altering cellular metabolism to drive anabolic processes necessary for cell growth, including biosynthesis of proteins, lipids and nucleic acids, and by inhibiting catabolic processes, such as autophagy (1-4). mTORC1 recruitment and activation occurs via intracellular sensing of amino acids at the lysosomal membrane, which in turn activates the Ras-related GTPase (Rag) heterodimers (5-7). On the other hand, extracellular growth factor stimulation of receptor tyrosine kinases (RTKs) transmit signals through the phosphoinositide-3-kinase/RAC-serine/threonine-protein kinase (PI3K/Akt) axis, resulting in the activation of mTORC1 via binding of the GTP-bound form of the small GTPase Ras homologue enriched in brain (Rheb) (1). This process critically involves the inhibition of the Rheb GAP tuberous sclerosis complex (TSC), formed of hamartin (TSC1), tuberin (TSC2), and a Tre2-Bub2-Cdc16 1 domain family, member 7 (TBC1D7) (8, 9). The TSC2 subunit contains the GAP domain, maintaining Rheb in the inactive GDP-bound form in the absence of growth signal stimuli. TSC2 associates with lysosomes but quickly relocalizes to the cytoplasm upon amino acid or growth factor stimulation (10, 11). This enables Rheb-GTP to bind mTOR and activate mTORC1, which propagates signals to downstream effectors such as ribosomal protein S6 (S6P) and eukaryotic translation initiation factor 4E-binding protein 1 (4EBP1) that promote protein synthesis and cell growth (12, 13). Although overall Rheb-GTP levels are high in cells (14, 15), a guanine nucleotide exchange factor (GEF) for Rheb remains elusive.

Rheb is a member of the large family of Ras GTPases and contains a highly conserved G-domain, which is critical to its function in signal transduction (16). Rheb, like all Ras proteins, is post-translationally modified by the addition of a farnesyl moiety via a stable thioether linkage to the Cys residue of its C-terminal CAAX motif (17-19). Additional targeting features such as reversible palmitoylation for N- and H-Ras and a polybasic stretch for K-Ras, upstream of the CAAX motif in the hypervariable region (HVR) maintain these Ras family proteins at the plasma membrane (PM) (17, 20-23). Due to the lack of these secondary targeting features, farnesylated Rheb associates with any membrane in the cell, causing its partitioning to the vast endomembrane surfaces of the cell (24, 25). However, in addition to this unspecific partitioning to cellular membranes, a significant enrichment on perinuclear membranes on and proximal to the late endosome/lysosome has been observed (7, 11, 26-28).

It was shown that Rheb interacts with the GSF PDEδ (delta subunit of phosphodiesterase-6) via its C-terminal farnesyl (24, 29, 30). Furthermore, PDEδ plays an essential role in spatial cycles that maintain prenylated Ras proteins on the PM by sequestering them in the cytosol (22, 24, 30). A small GTPase ADP-ribosylation factor like 2 (Arl2) mediates the release of farnesylated Ras from PDEδ on perinuclear membranes in a GTP dependent manner (22, 30). This concentrates Ras on perinuclear membranes, where electrostatic interaction traps K-Ras on the recycling endosome, and palmitoyl addition via palmitoyltransferase (PAT) traps the H- and N-Ras on the Golgi apparatus. Association with the anterograde vesicular transport from these organelles then reinstates PM localization of Ras proteins (31). Rheb lacks the secondary targeting features that enable anterograde transport to the PM. Therefore, we hypothesized that Arl2-mediated release from PDEδ concentrates Rheb on perinuclear lysosomal/late endosomal membranes where its effector mTOR resides. Herein, we do not only show that Rheb localization is indeed maintained by an energy-driven PDEδ-Arl2 mediated spatial cycle, but that this cycle also drives its GTPase cycle, which is essential to maintain mTORC1 responsiveness to growth factors.

## RESULTS

### Growth factor stimulation affects the partitioning of Rheb between perinuclear membranes and the cytosol

To determine the subcellular localization of Rheb, we compared the distribution of ectopically expressed Rheb, N-terminally tagged with mCitrine (mCitrine-Rheb) with the endogenous localization of Rheb, as determined by immunofluorescence (IF) using a Rheb specific antibody in mouse embryonic fibroblasts (MEFs) immortalized through a p53 knockout (TSC2+/+ MEFs from here on). Both proteins were found on endomembranes, with a significant enrichment on perinuclear membranes that coincided with the localization of the Rheb effector mTOR (**Fig. 1a**), as well as with the late endosome/lysosome marker, Rab 7 (**Fig. 1b**). However, a significant fraction of Rheb could also be observed in the cytosol.

**Fig. 1.**
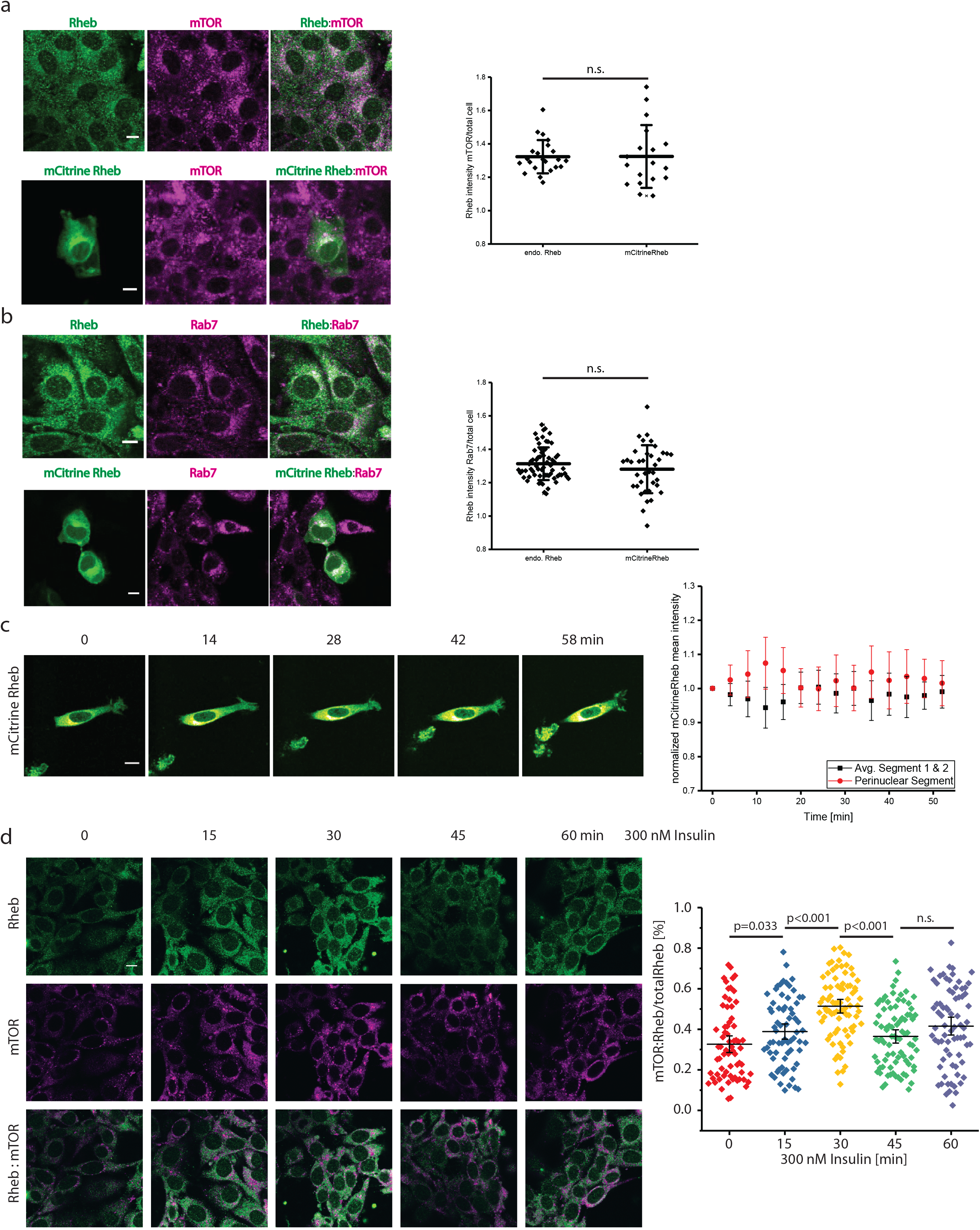
Rheb partitioning to perinuclear membranes is enhanced by insulin. (**a,b**) Confocal micrographs of TSC2+/+ MEFs depicting the spatial distribution of immunofluorescence signal for mTOR (**a**) and Rab7 (**b**) with endogenous Rheb (upper row) or mCitrine-Rheb (lower row). Graphs: Ratio of Rheb or mCitrine-Rheb co-localizing with mTOR (**a**) or Rab7 (**b**) over total cell Rheb or mCitrine-Rheb for individual cells with mean fluorescence intensity ± S.D. (n>20 cells per condition from two independent experiments). P-values were obtained by Student’s t-test, >0.05 labeled as not significant (n.s.). (**c**) Time series of serum-starved TSC2+/+ MEFs expressing mCitrine-Rheb treated with 300 nM insulin. Rheb spatial distribution was determined by segmenting cells into 3 segments of equal radius from the plasma membrane (PM) to the nuclear membrane (NM) (1-3) and calculating the mean fluorescence intensity in each segment normalized to the total intensity in all three segments (see Methods). Black: normalized average of 1^st^ and 2^nd^ segment; red: normalized average of perinuclear 3^rd^ segment. Error bars depict s.d.; n=7 cells from two independent experiments. (**d**) Left: Confocal micrographs of serum-starved TSC2+/+ MEFs depicting localization of immunofluorescence signal for endogenous Rheb (first row) and mTOR (second row) at different time points after 300 nM insulin stimulation. Third row: overlaid images. Right: Fraction of Rheb mean fluorescence intensity co-localizing with mTOR fluorescence in single cells (n>67 for each condition from two independent experiments). Line depicts mean and error bars depict 95% confidence interval. P-values were obtained by one-way ANOVA. Scale bars: 10 μm.

To examine whether the partitioning of Rheb between cytosol and membranes is affected by the activation of mTOR upon growth factor stimulation, we treated serum-starved TSC2+/+ MEFs with insulin and monitored the localization of mCitrine-Rheb over time by radial segmentation of the cells into 3 spatial bins from plasma membrane (PM) to nuclear membrane (NM) within a 60° angle around the mCitrine-Rheb intensity-weighted longitudinal cellular axis. Upon insulin stimulation, mCitrine-Rheb was transiently recruited to perinuclear membranes of the cell, suggesting the local interaction of active Rheb-GTP with the effector mTOR (**Fig. 1c**). Consistent with this, IF for endogenous Rheb and mTOR at different time points after insulin stimulation of serum-starved TSC2+/+ MEFs also displayed a transient recruitment of endogenous Rheb to mTOR containing membranes (**Fig. 1d**). Enrichment of endogenous Rheb on mTOR-rich membranes was slightly delayed compared to ectopically expressed mCitrine-Rheb, (30 min in the IF versus 12 min in the live-cell time course), reflecting a shift in the reaction rate upon increasing the concentration of a reactant according to the law of mass action. This transient increase in Rheb-enrichment on mTOR containing membrane indicates that the interchange between the soluble and membrane-bound fraction of Rheb is dynamic, likely mediated and maintained by the solubilizing factor PDEδ.

### Arl2-mediated localized release from PDEδ deposits Rheb on perinuclear membranes

We examined how the interaction of Rheb with PDEδ influences its cytosol-membrane partitioning by inhibiting PDEδ function using the small-molecule inhibitor Deltarasin that targets PDEδ’s prenyl-binding pocket (29). For this, we monitored the localization of Rheb, N-terminally tagged with mCherry (mCherry-Rheb) as well as the hypervariable region of Rheb, N-terminally tagged with mCitrine (mCitrine-Rheb HVR) in TSC2+/+ MEFs after treatment with 3 μM Deltarasin. This resulted in a gradual loss of the perinuclear enrichment for both proteins, demonstrating that the interaction of PDEδ via the farnesyl group in the HVR is necessary for generating the perinuclear concentration of Rheb (**Fig. 2a**). We also directly monitored the effect of Deltarasin on the interaction between Rheb and PDEδ by fluorescence lifetime imaging microscopy of Förster resonance energy transfer (FLIM-FRET), with mCitrine-Rheb as the donor and PDEδ, C-terminally tagged with mCherry (mCherry-PDEδ) as FRET acceptor (29, 32, 33) (**Fig. 2b**). The interaction between mCitrine-Rheb and mCherry-PDEδ was apparent from the decrease in donor fluorescence lifetime of mCitrine-Rheb upon co-expression of mCherry-PDEδ (**Fig. 2c**). In order to quantify how mCherry-PDEδ expression affects this interaction, we computed the fraction (α) of mCitrine-Rheb interacting with mCherry-PDEδ from the fluorescence decay profiles by global analysis (34) (**Fig. 2b,d**). α increased with the expression level of mCherry-PDEδ, reflecting that the amount of detected mCitrine-Rheb/mCherry-PDEδ complexes is limited by the availability of the solubilizer, mCherry-PDEδ (**Fig. 2d**). Treatment with Deltarasin clearly diminished the interaction that occurs throughout the cytoplasm and led to an increased overall membrane deposition of mCitrine-Rheb as indicated by the loss of nuclear mCitrine-Rheb fluorescence (**Fig. 2e**). These experiments indicate that while the interaction of Rheb with PDEδ causes a fraction of Rheb to be partitioned to the cytosol, it also drives the perinuclear enrichment of Rheb, countering equilibration to other endomembranes. We therefore hypothesized that activity of Arl2 at perinuclear membranes causes local deposition of Rheb by releasing it from PDEδ.

**Fig. 2.**
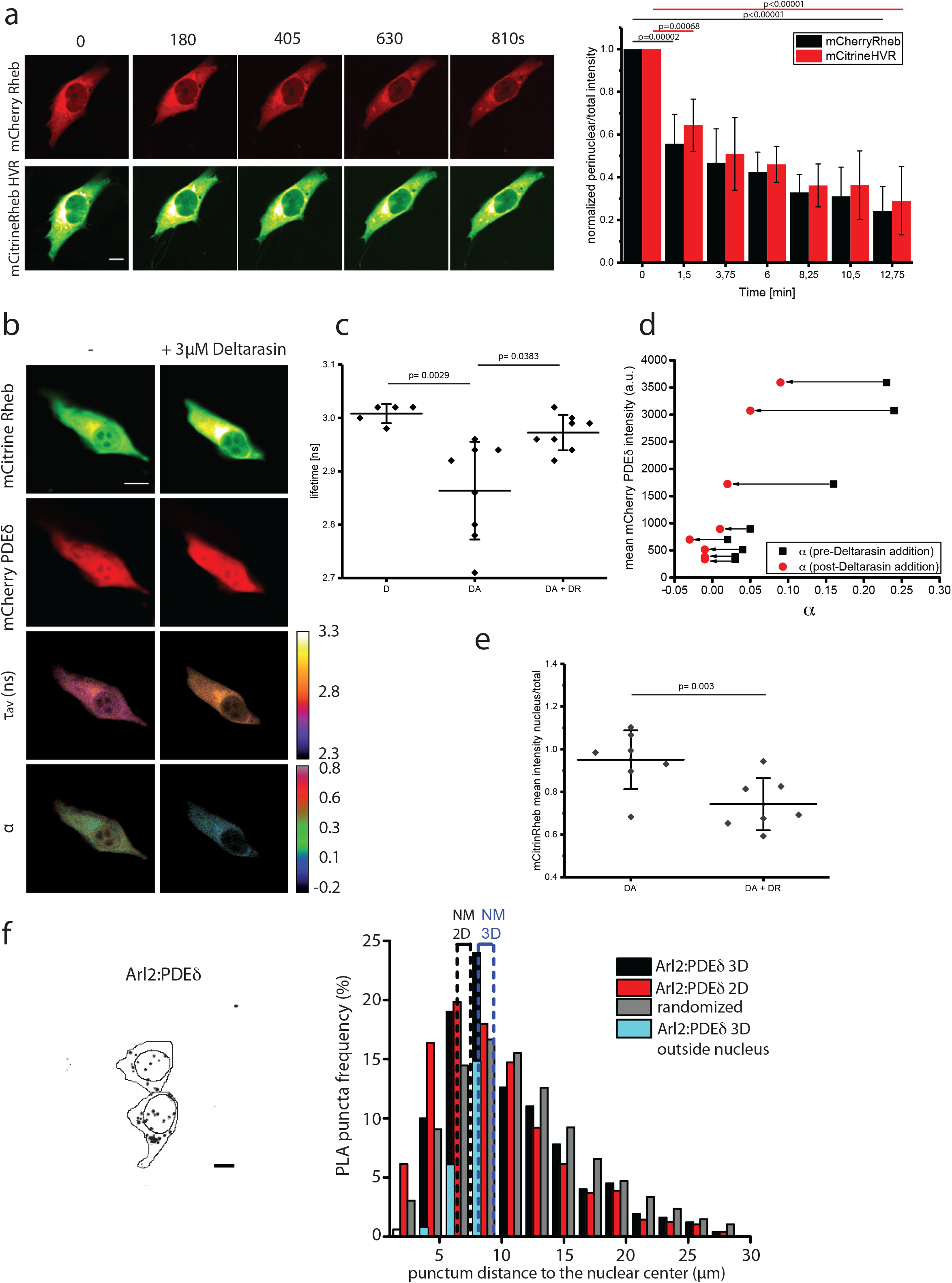
PDEδ interacts with Rheb in the cytoplasm and Arl2 on perinuclear membranes. (**a**) Time series of TSC2+/+ MEFs co-expressing mCherry-Rheb (upper row) and mCitrine-Rheb-HVR (lower row), treated with 3μM Deltarasin at t=0 s. Right plot: normalized perinuclear over total fluorescence of mCherry-Rheb (black) and mCitrine-Rheb-HVR (red) over time (N=4). (**b-e**) FRET-FLIM measurements of mCitrine-Rheb and mCherry-PDEδ interaction in TSC2+/+ MEFs, before (-) and 30 min after addition of 3 μM Deltarasin. (**b**) Fluorescence intensity of the indicated proteins (upper two rows), spatial distribution of the mean fluorescence lifetime (τ_av_) (third row) and the molar fraction of interacting molecules (α) (fourth row). **(c)** τ_av_ of donor only (D), donor-acceptor complex (DA) and donor-acceptor complex after Deltarasin addition (DA+DR) in nanoseconds (n=8 cells). **(d)** Change in α after Deltarasin addition as function of mCherry-PDEδ expression in individual cells. Black arrows: decrease in α after Deltarasin addition. **(e)** Graph depicting the ratio of mean nuclear over total cell mCitrine-Rheb intensity before (DA) and after Deltarasin addition (DA+DR). **(f)** Representative maximum intensity projection of a confocal Z-stack of Arl2-PDEδ PLA fluorescent interaction puncta (left) from TSC2+/+ MEFs. The cell outline was determined from the integrated fluorescence images of the stack, while nuclei were identified with DAPI staining. Histogram (right) depicts the mean distribution of distances of Arl2/PDEδ PLA puncta to the nuclear centre obtained from 3D cell volumes (black) and corresponding 2D maximal intensity projections (red). For comparison, the mean distance distribution of all pixels within cells (grey) is shown. Dotted lines represent the average position of the nuclear membrane (NM) for both 3D (blue) and 2D-projections (black). Puncta detected in 3D as positioned outside of the nucleus but within the nuclear/perinuclear segments are represented in turquoise (n= 33 cells, data from 2 independent experiments). P-values obtained by Student’ s t-test. Scale bars: 10 μm. All data are mean±S.D.

The small GTPases Arl2 and Arl3 bind PDEδ in a GTP-dependent manner, thereby inducing an allosteric release of farnesylated cargo from PDEδ (30, 35). Only Arl2 was shown to be essential for the maintenance of the PM enrichment of K-Ras (22) and an Arl2 GEF has so far not been identified. The subcellular locus of allosteric Arl2 release activity on PDEδ has however been demonstrated to occur in a region on or proximal to the recycling endosome (22). IF using an Arl2 specific antibody in TSC2+/+ MEFs showed that Arl2 proteins reside on perinuclear membranes, as well as in the nucleus and to a lesser extent in the cytoplasm (Supplementary **Fig. 1a**). This indicates that Arl2 proteins partition between the cytosol and membranes, consistent with biophysical data that shows that Arl proteins interact with membranes via their N-terminal amphipathic helices (36). In comparison, IF with a PDEδ specific antibody only showed nuclear and diffuse cytoplasmic staining, consistent with the soluble state of the protein.

To investigate if and where Arl2-mediated release takes place, we measured the interaction between endogenous PDEδ and Arl2 by in situ proximity ligation assay (PLA) (37). The PLA reaction generates discrete fluorescent puncta in areas of the cell where protein interactions occur (38). To obtain information on the radial distribution of this interaction, we computed the distance for each PLA punctum to the center of the nucleus in many cells (see ‘Methods’, **Fig. 2f, Supplementary Fig. 1b**). The puncta distributions for Arl2/PDEδ peaked at the position of the nuclear membrane and the adjacent nuclear spatial bin, to rapidly decay towards the cell periphery. However, the shape of these puncta distributions is biased by the cell shape and size (**Fig. 2f**, ‘Methods’). To correct for this bias in the puncta distributions, a pixel-distance distribution to the center of the nucleus that reflects the cell shapes was subtracted (**Supplementary Fig. 1b**). The positive peak around the average position of the nuclear membrane and the negative broad peak in the cytoplasmic area in these distance distributions, show that the interaction between PDEδ and Arl2 predominantly occurs near perinuclear membranes of the cell, which implies that allosteric release of Rheb from PDEδ occurs in this area. Additionally, the 3D distribution of each PLA punctum to the nuclear center showed that the majority of the puncta in the spatial bins proximal to the nuclear membrane were located outside of the nucleus. This confirmed the results obtained from the 2D projections that the allosteric release of Rheb from PDEδ via Arl2 activity occurs in the perinuclear area of the cell.

The transient increase in Rheb localization in the perinuclear area upon insulin stimulation (**Fig. 1c,d**) suggests that growth factor stimulation can either cause increased deposition on or increased retention of Rheb at perinuclear membranes. The Rheb-GAP TSC2 dissociates from lysosomes to the cytoplasm upon growth factor stimulation (11), thereby likely causing an increase in Rheb-GTP on lysosomal membranes that can bind and activate mTOR. To evaluate whether TSC2, and thereby the activation state of Rheb, can influence its steady state localization, we compared the perinuclear enrichment of mCitrine-Rheb in TSC2+/+ MEFs to that in TSC2-/- MEFs, an isogenic cell line in which TSC2 is knocked out, resulting in constitutively active Rheb-GTP (39, 40). The perinuclear membrane enrichment of mCitrine-Rheb was significantly increased in the TSC2-/- MEFs as compared to the TSC2+/+ MEFs (**Supplementary Fig. 2**) indicating that the phosphorylation state of the guanine nucleotide bound to Rheb increases its retention on perinuclear membranes.

To investigate how the PDEδ mediated partitioning of Rheb is coupled to its GTPase cycle, we generated PDEδ knockouts of TSC2+/+ and TSC2-/- MEFs by CRISPR-Cas9 (41, 42) using a single guide RNA targeting PDEδ (TSC2+/+ sgRNA PDEδ) and an empty Cas9 vector (TSC2+/+ E.V.) as control. Evaluation of perinuclear enrichment of mCitrine-Rheb in these cells by segment analysis (**Fig. 3a,b**) revealed a significant decrease for the TSC2+/+ sgRNA PDEδ cells, as well as a concurrent increase in fluorescence intensity in the periphery of the cell (**Fig. 3a**). This is consistent with the experiments shown in **Fig. 2a,b** that demonstrate that Rheb solubilization by PDEδ is necessary to maintain a concentration of Rheb on perinuclear membrane compartments. In contrast, the radial intensity profile of mCitrine-Rheb in TSC2-/- MEFs was unaffected by PDEδ knockout (**Fig. 3b**). This indicates that in absence of TSC2 perinuclear Rheb enrichment is determined by its interaction with mTOR, uncoupling its enrichment from PDEδ-mediated deposition.

**Fig. 3.**
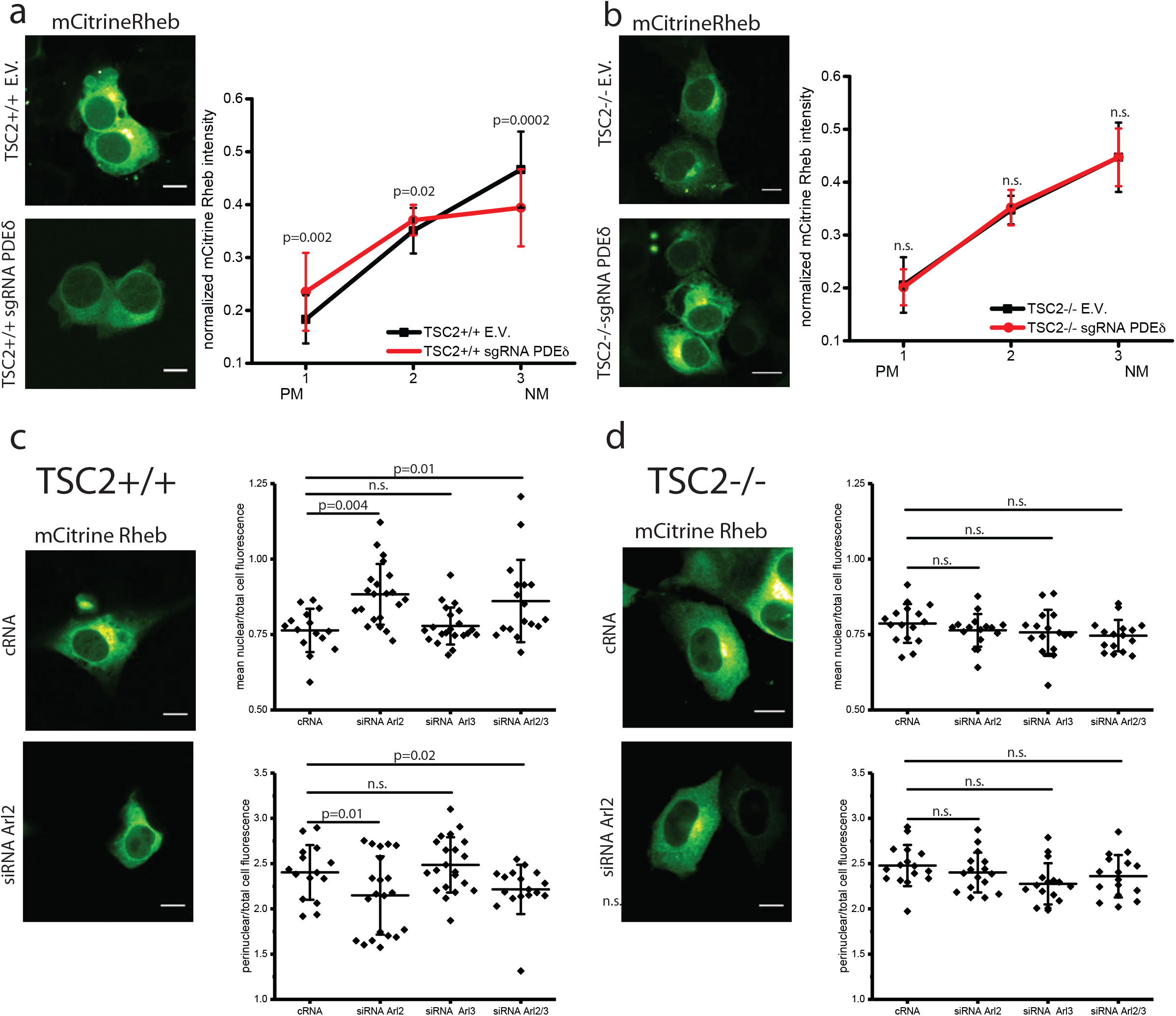
The PDEδ/Arl2 system maintains Rheb localization. Localization distribution of Rheb variants in TSC2+/+ MEFs (**a**) and TSC2-/- MEFs (**b**) stably expressing either an empty Cas9 vector (E.V.) or a Cas9 vector encoding a single guide RNA for silencing PDEδ (sgRNA PDEδ), ectopically expressing mCitrine-Rheb (**a,b**). The steady state localization of mCitrine-Rheb was determined by segment analysis with angular masking. Radial profiles depict the normalized fluorescence intensity in each segment (n> 15 cells for each cell line from two independent experiments). (**c,d**) Dependence of Rheb localization on Arl2/3 in TSC2+/+ (c) and TSC2-/- MEFs (**d**). Representative maximum intensity projections (left micrographs) of confocal Z-stacks of MEFs transfected with Arl2 targeting siRNA (siRNA Arl2) or non-targeting siRNA control oligonucleotides (cRNA), ectopically expressing mCitrine-Rheb. Dot plots (right) depict the ratio of mean nuclear over total cell mCitrine-Rheb intensity (upper graphs) or the ratio of mean perinuclear over total cell mCitrine-Rheb intensity (lower graphs) for individual cells harboring control RNA (cRNA), siRNA Arl2, siRNA Arl3 and double knockdown (siRNA Arl2/3) (n> 16 cells for each condition from 2 independent experiments). Data is represented as mean±S.D.. P values were obtained by Student’s t-test, >0.05 are labeled as not significant (n.s.). Scale bars: 10μm.

If Arl2 acts as the factor releasing Rheb from PDEδ in the perinuclear area to maintain Rheb enrichment there, siRNA-mediated knockdown of Arl2 should disrupt perinuclear enrichment of Rheb and increase the fraction of PDEδ-bound, soluble Rheb. Indeed, in TSC2+/+ cells, Arl2 knockdown resulted in a significant decrease in perinuclear enrichment of mCitrine-Rheb and an increased soluble fraction of mCitrine-Rheb, as apparent from its increased nuclear intensity (**Fig. 3c, graphs**). This shows that Arl2 activity unloads Rheb from PDEδ onto perinuclear membranes. In contrast, Arl2 knockdown had no effect on mCitrine-Rheb solubilization or perinuclear enrichment in TSC2-/- MEFs (**Fig. 3d, graphs**), providing further evidence that in absence of TSC2, Rheb-GTP is stably associated with perinuclear membranes due to interaction with its effector mTOR. Arl3 knockdown had no effect on mCitrine-Rheb localization, neither in TSC2+/+ nor TSC2-/- MEFs (**Fig. 3c,d**), showing that Arl3 does not allosterically release Rheb from PDEδ. These results show that perinuclear membrane localized Arl2-GTP activity mediates release of Rheb from PDEδ onto perinuclear membranes. This release mechanism thereby generates a directional flux in Rheb cycling between membranes and cytosol, where GTP hydrolysis happens on TSC2-containing perinuclear membranes.

To further substantiate this, we investigated the radial distribution of mCitrine-Rheb-HVR, which interacts with PDEδ via the farnesyl tail but cannot interact with mTOR due to the lack of the Rheb G-domain. In both TSC2+/+ and TSC2-/- sgRNA PDEδ MEFs, the distribution of mCitrine-Rheb HVR decreased significantly towards the perinuclear segment of the cells as compared to their E.V. controls (**Fig. 4a**) showing that RhebGTP retention on perinuclear membranes occurs via mTOR interaction. In full agreement with this notion, the radial profiles of the active EYFP-Rheb Q64L mutant, which was shown to display a higher GTP loading status than wild type Rheb (14, 43), were independent of PDEδ in both examined cell types (**Fig. 4b**). These experiments indicate that interference with the hydrolysis of GTP-loaded Rheb either by knock-out of TSC2 or by a constitutively active mutant enables a sufficiently strong level of Rheb-effector interaction to retain Rheb on perinuclear membranes without requiring continuous re-deposition by PDEδ. To substantiate that Rheb-GTP interacting with effectors on membranes is poorly re-solubilized by PDEδ, we quantified the perinuclear enrichment of mCitrine-Rheb upon ectopic mCherry-PDEδ expression in TSC2+/+ and TSC2-/- cells (**Fig. 4c**). In TSC2+/+ MEFs perinuclear enrichment of mCitrine-Rheb was decreased upon ectopic mCherry-PDEδ expression. In contrast, mCitrine-Rheb perinuclear enrichment was unaffected by ectopic PDEδ expression in TSC2-/- cells. These results indicate that perinuclear membrane associated Rheb-GTP cannot be (re)-solubilized by PDEδ due to its interaction with effectors on membranes.

**Fig. 4:**
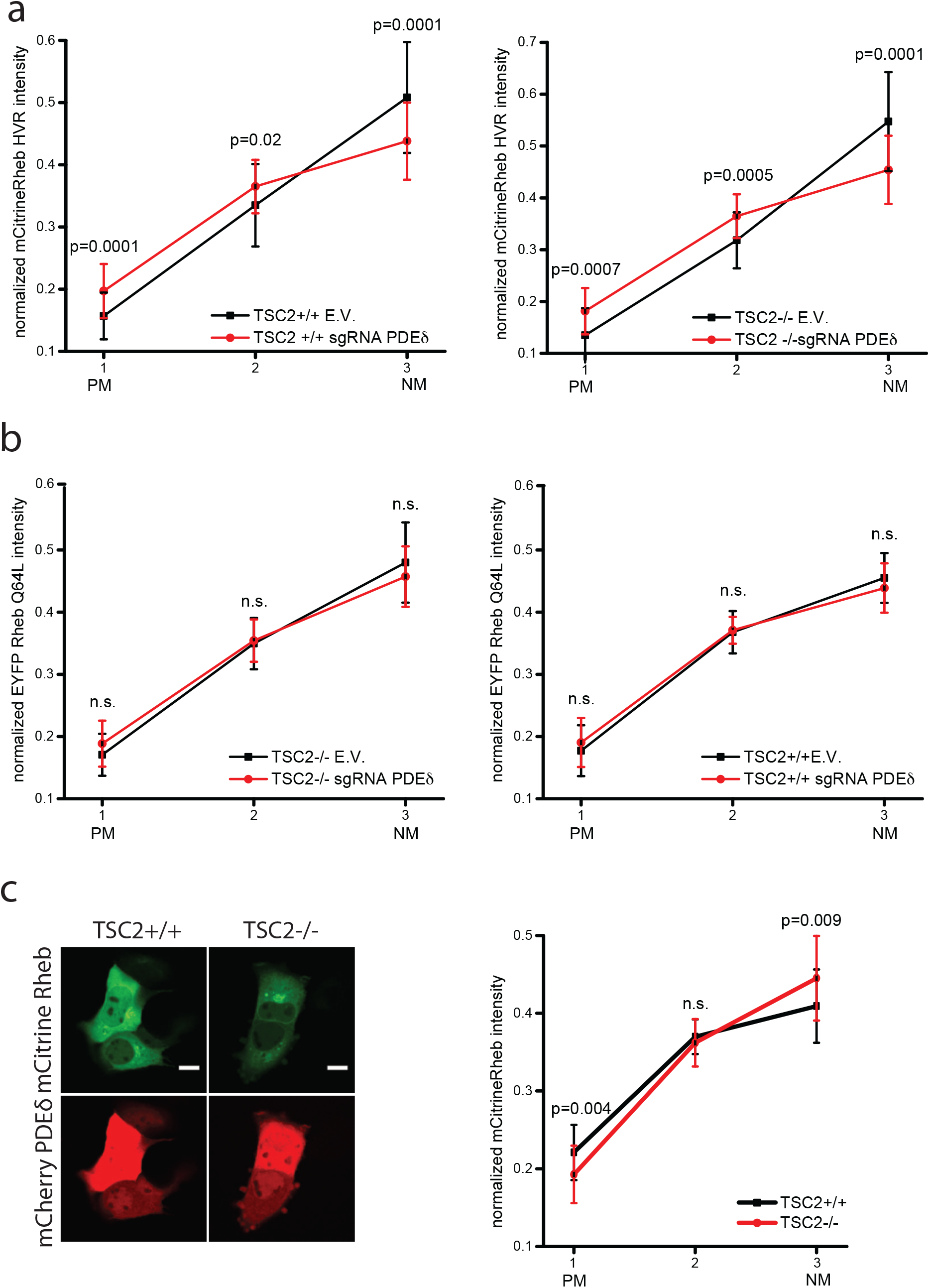
GTP-loaded Rheb is retained on perinuclear membranes. Localization of Rheb variants in TSC2+/+ MEFs (left) and TSC2-/- MEFs (right) stably expressing either an empty Cas9 vector (E.V.) or a Cas9 vector encoding a single guide RNA for silencing PDEδ (sgRNA PDEδ), transiently expressing either mCitrine-Rheb HVR (**a**) or the constitutively active EYFP-Rheb Q64L mutant (**b**). The steady state localization of the proteins was determined by segment analysis with angular masking. Radial profiles depict the normalized fluorescence intensity for each transiently expressed protein in each segment ±S.D. for the indicated cell line (n> 15 cells for each condition from two independent experiments). (**c**) Confocal micrographs of TSC2+/+ MEFs (left column) and TSC2-/- MEFs (right column) co-expressing mCitrine-Rheb (upper row) and mCherry-PDEδ (lower row). Steady state localization of mCitrine-Rheb was quantified by segment analysis. Radial profiles as in **a,b** (n> 20 cells for each condition from two independent experiments, TSC2+/+: black, TSC2-/-: red). P values were obtained by Student’s t-test, >0.05 are labeled as not significant (n.s.). Scale bars: 10 μm.

### mTORC1 signaling depends on PDEδ and Arl2

To investigate the functional implications of PDEδ-mediated solubilization of Rheb on its growth factor-controlled GTPase cycle, we compared the effect of PDEδ knockout on mTORC1 signaling in TSC2+/+ and TSC2-/- cells. The signaling output of mTORC1 was determined by the level of ribosomal protein S6 (S6P) phosphorylation in response to 300 nM insulin for 15 minutes. PDEδ knockout significantly reduced both, basal and insulin-induced S6P phosphorylation in TSC2+/+ MEFs (**Fig. 5a, Supplementary Fig. 3**). In contrast, PDEδ knockout in TSC2-/- MEFs showed no effect on basal S6P phosphorylation that was independent on insulin stimulation (**Fig. 5b, Supplementary Fig. 4**). In agreement with these results, siRNA-mediated knockdown of Arl2 and not Arl3 led to a decrease in S6P phosphorylation only in TSC2+/+ MEFs, while in TSC2-/- MEFs phosphorylation of S6 remained unchanged (**Fig. 5c,d, Supplementary Fig. 5**).

**Fig. 5:**
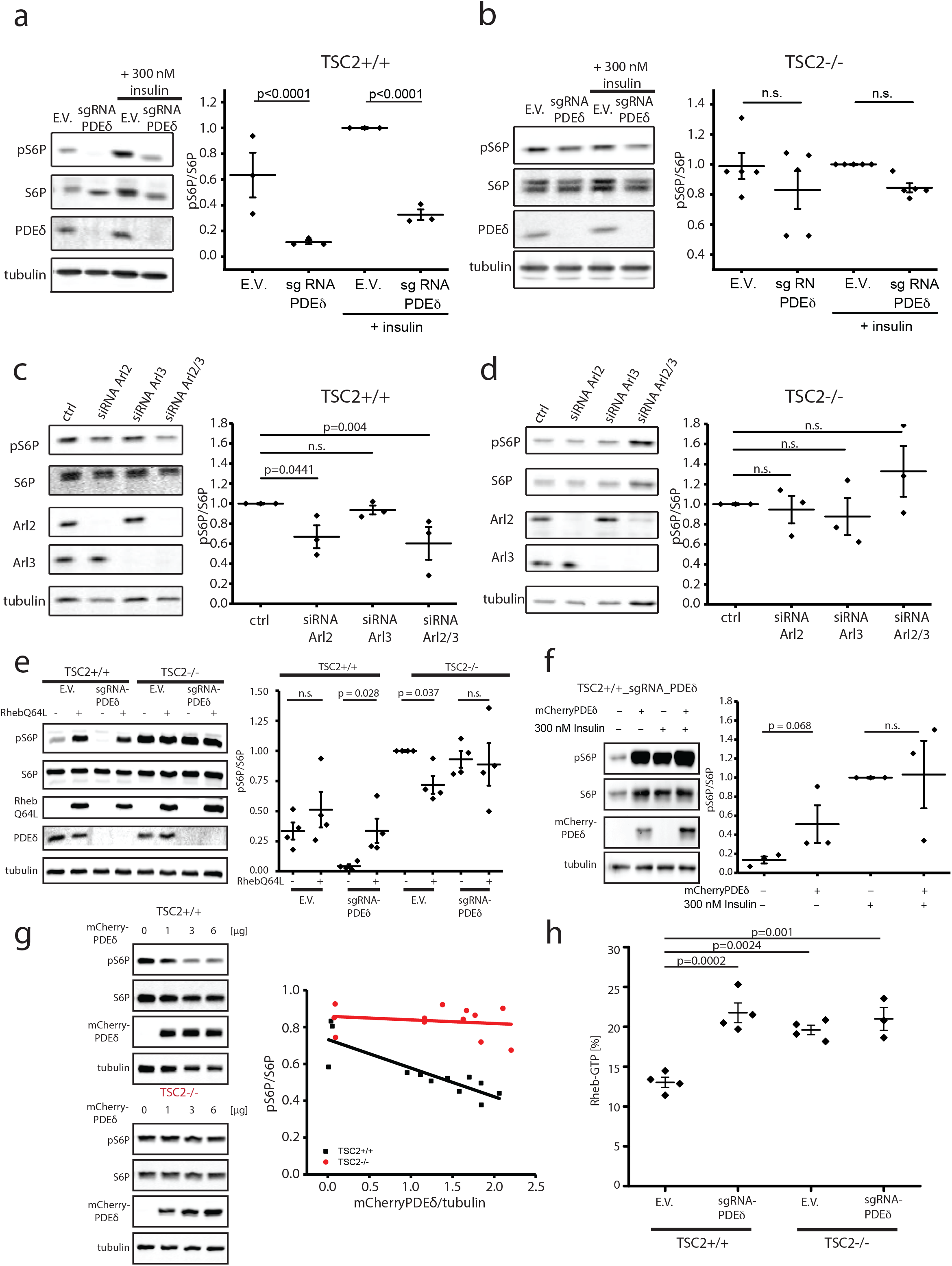
The PDEδ/Arl2 system mediates Rheb-dependent mTORC1 signaling. (**a,b**) S6P phosphorylation in serum-starved TSC2+/+ E.V. and TSC2+/+ sgRNA PDEδ MEF (**a**) and TSC2-/- E.V. and TSC2-/- sgRNA PDEδ MEF (**b**) prior and post insulin stimulation. Representative western blots (left) show phosphorylation of S6P (pS6P), total levels of S6P (S6P), PDEδ and tubulin (loading control). Dot plots (right) depict the level of pS6P/S6P normalized to the insulin-stimulated control cell line (E.V. + insulin) (TSC2+/+: n=3; TSC2-/-: N=5) for each individual blot. (**c,d**) S6P phosphorylation in TSC2+/+ (c) and TSC2-/- MEFs (**d**) upon siRNA-mediated knockdown of Arl2, Arl3 or both determined by western blot 48h post siRNA transfection. Representative blots (left) show phosphorylation of S6P (pS6P), total levels of S6P (S6P), Arl2, Arl3 and tubulin (loading control). Dot plots (right) depict the level of pS6P/S6P normalized to control (non-targeting siRNA) (N=3) for each individual blot. P values >0.05 are labeled as not significant (n.s.). (**e,f**) S6P phosphorylation in serum-starved TSC2+/+ and TSC2-/- MEFs with E.V. or sgRNA PDEδ and with/without transient mCitrine-RhebQ64L (**e**) or serum-starved TSC2+/+ sgRNA PDEδ with/without mCherry-PDEδ (**f**) expression. S6P phosphorylation levels in (**f**) were determined prior and post-insulin stimulation. Representative western blots (left) show phosphorylation of S6P (pS6P), total levels of S6P (S6P), mCitrineRhebQ64L or mCherryPDEδ expression and tubulin (loading control). Dot plots (right) depict the level of pS6P/S6P normalized to TSC2-/- E.V. for each individual blot (**e**) or insulin-stimulated TSC2+/+ sgRNA PDEδ (**f**) (N=4 and 3, respectively). (**g**) Dependence of S6P phosphorylation on ectopic mCherry-PDEδ expression (24hrs post-transfection) in TSC2+/+ (black) and TSC2-/- MEFs (red). Representative western blots (left) show phosphorylation of S6P (pS6P), total levels of S6P (S6P), mCherryPDEδ and tubulin (loading control). Scatter plot (right) shows the relative level of pS6P/S6P as function of mCherry-PDEδ expression normalized to the tubulin loading control. Lines represent linear fits to the data (N=3). P values were obtained by Student’s t-test, >0.1 are labeled as not significant (n.s.) (**h**) Level of Rheb-GTP (GTP/(GTP+GDP)) in TSC2+/+ and TSC2-/- MEFs with E.V. or sgRNA PDEδ determined by thin layer chromatography of radioactive GTP/GDP from immunoprecipitated endogenous Rheb (N= 4). P values were obtained by One-way ANOVA using Turkey’s multiple comparison. Insulin stimulation: 300nM for 15 minutes. Error bars depict S.E.M.

To investigate whether increased GTP loading of Rheb could rescue the mTOR signaling deficit induced by PDEδ knockout, we ectopically expressed the EYFP-Rheb Q64L mutant. Indeed, in TSC2+/+ sgRNA PDEδ, RhebQ64L expression raised S6P phosphorylation back to a level comparable to TSC2+/+ E.V. cells, while a slight increase in S6P phosphorylation was observable in TSC2+/+ E.V.. In contrast, RhebQ64L expression did not significantly alter the higher basal S6P phosphorylation levels in TSC2-/- MEF cells with or without PDEδ knockout (**Fig. 5e, Supplementary Fig. 6**).

To confirm that the loss of S6P phosphorylation is indeed a consequence of the PDEδ knockout, we transiently expressed a mCherry-PDEδ construct in TSC2+/+ MEF cells with and without PDEδ knockout and studied basal and insulin stimulated S6P phosphorylation. As expected, PDEδ re-expression led to an increase of pS6P in the TSC2+/+ sgRNA PDEδ MEFs while in the TSC2+/+ E.V. ectopic PDEδ expression had no significant effect on the pS6P level (**Fig. 5f, Supplementary Fig. 7**). Complementary to this, increasing mCherry-PDEδ expression in parental TSC2+/+ MEFs decreased S6P phosphorylation, while mCherry-PDEδ expression had no effect on S6P phosphorylation in the TSC2-/- MEFs (**Fig. 5g, Supplementary Fig. 8**). This indicates that there is an optimal PDEδ concentration that enables S6P signaling: Both, a too low and a too high level of PDEδ render Rheb-GTP enrichment on perinuclear membranes insufficient to allow robust S6P signaling. In the former case, too few PDEδ molecules are available to solubilize Rheb molecules from endomembranes and the low concentration of PDEδ loaded with Rheb becomes rate limiting in the Arl mediated release in the perinuclear area. In the latter case (high PDEδ concentration), the fraction of PDEδ loaded with Rheb becomes so low with respect to total PDEδ that the Arl-mediated release operates mostly on PDEδ without cargo, thereby generating a futile cycle.

These experiments demonstrate that PDEδ-mediated solubilization of Rheb is essential for activating mTORC1 signaling in response to growth factor signals. The retention of Rheb-GTP at perinuclear membranes by interaction with its effector mTOR implies that the Rheb species dissociating from perinuclear membranes before solubilization by PDEδ must primarily be Rheb-GDP and indicates that nucleotide exchange happens on soluble, PDEδ-bound Rheb. Since Rheb was speculated to function independent of a GEF and was shown to exhibit a significantly higher intrinsic nucleotide exchange rate in solution versus membrane (44), we hypothesized that the dynamics of the PDEδ/Arl2 system could generate a constant re-flux of Rheb-GTP to perinuclear membranes, where the growth-factor regulated level of TSC2 activity determines the level of S6P phosphorylation. To investigate this, we labeled cells using radioactive ortho-phosphate, immuno-precipitated Rheb from the lysates and separated GTP and GDP fractions by thin layer chromatography. TSC2+/+ sgRNA PDEδ exhibited a significantly higher GTP/GDP ratio compared to TSC2+/+ E.V., while there was no difference between TSC2 -/- E.V. and TSC2-/- sgRNA PDEδ GTP/GDP ratios. Moreover, the TSC2+/+ sgRNA PDEδ GTP/GDP ratio resembled that of TSC2 knock out cell lines. The increased Rheb GTP level upon PDEδ knockout (**Fig. 5h, Supplementary Fig. 9**) in conjunction with the loss of perinuclear Rheb to other endomembranes (**Fig. 3a**) and the dramatic reduction in S6P phosphorylation under the same condition (**Fig. 5a**), implies that the PDEδ-Arl2 localization system is indeed essential to promote the reflux of Rheb that has undergone nucleotide exchange in the cytoplasm to perinuclear membranes, where it is inactivated by TSC2 in absence of growth factor signals.

### Cell growth depends on the PDEδ-mediated spatial cycle of Rheb

The dependence of growth factor induced mTORC1 activity on PDEδ suggested that disruption of PDEδ-mediated solubilization of Rheb will have an inhibitory effect on cell growth only in TSC2+/+ cells that are responsive to growth factors. We therefore monitored the effect of PDEδ knockout on cell growth by real time cell analyzer (RTCA) and colony formation by clonogenic assays in TSC2+/+ as well as TSC2-/- MEFs.

PDEδ knockout resulted in a dramatic decrease of cell growth in TSC2+/+ MEFs, as compared to both E.V. and parental control cells, as apparent from RTCA growth curves as well as clonogenic assays (**Fig. 6a,c**). This is consistent with PDEδ-mediated solubilization of Rheb being necessary for mTORC1 responsiveness to growth factors. In contrast, no reduction in growth was observed in TSC2-/- PDEδ sgRNA MEFs, consistent with Rheb-GTP being uncoupled from the PDEδ-mediated solubilization cycle in these cells (**Fig. 6b,d**). Instead, a slightly increased growth rate was observed, likely reflecting that most Rheb-GTP is partitioned to membranes and therefore drives mTOR activation on lysosomes.

**Fig. 6.**
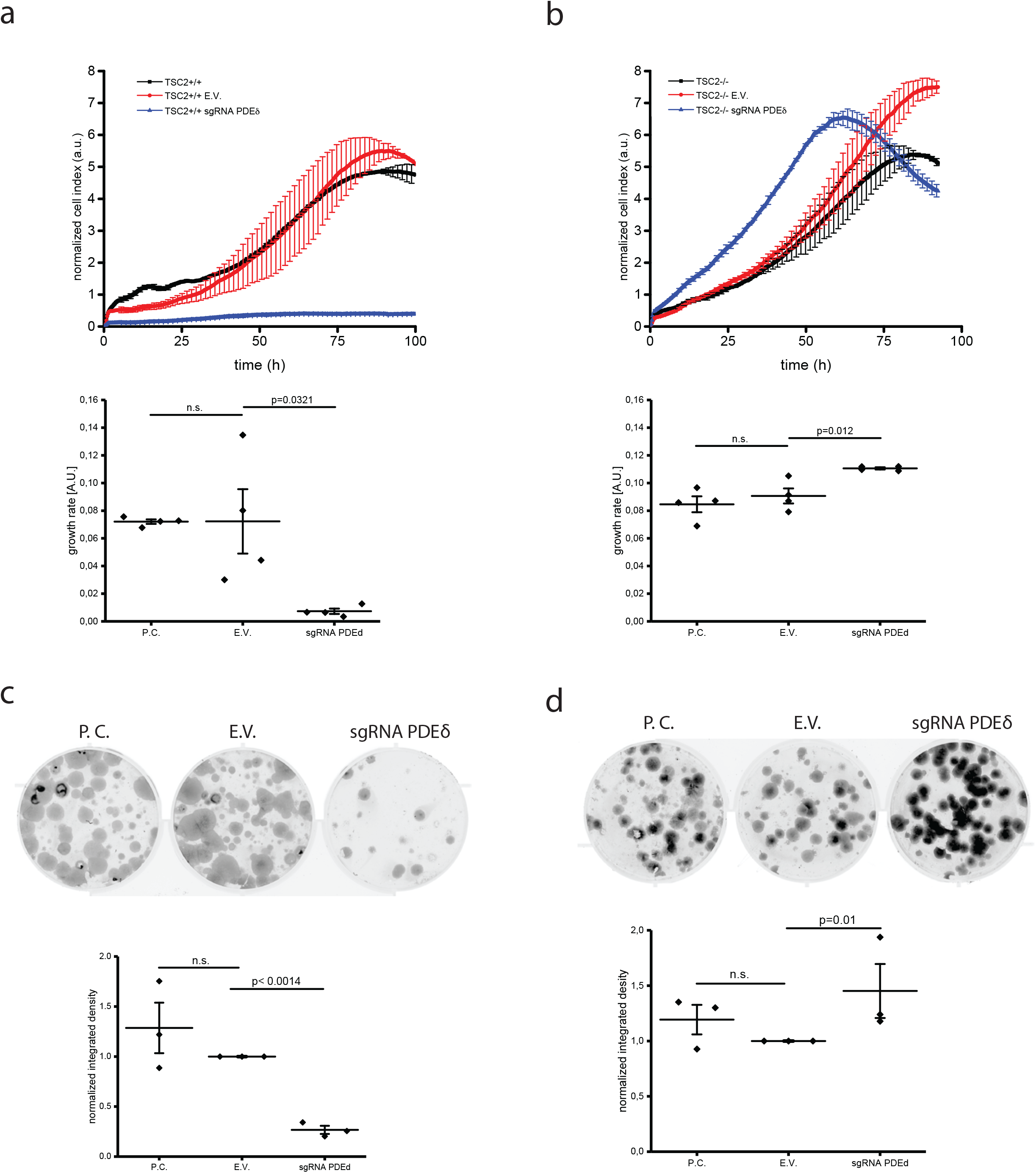
Cell growth depends on PDEδ in MEFs with regulated TSC2 activity. (**a,b**) Representative RTCA growth profiles for TSC2+/+ (**a**) and TSC2-/- MEF (**b**) parental cell lines (black trace) or stably expressing either an empty Cas9 vector (E.V; red trace) or a Cas9 vector encoding a single guide RNA for silencing PDEδ (sgRNA PDEδ; blue trace). Dot plots underneath depict the average growth rate ± S.E.M. between 40 and 60 hours (N=4 independent experiments in duplicate for all cell lines). (**c,d**) Clonogenic assays of TSC2+/+ MEF (**c**) and TSC2-/- MEF (**d**) parental cell lines (P.C.) or stably expressing either an empty Ca9 vector (E.V) or a Cas9 vector encoding a single guide RNA for silencing PDEδ (sgRNA PDEδ). Dot plots underneath depict the mean colony area coverage normalised to the E.V. control well (N=3 independent experiments, data are mean±S.E.M.). P values were obtained by Student’s t-test, >0.05 are labeled as not significant (n.s.).

## DISCUSSION

Here, we show that growth factor induced Rheb activity on mTORC1 is critically dependent on the solubilizing activity of PDEδ. This GSF causes the partitioning of Rheb between membranes and the cytosol. Arl2-GTP mediated localized release of Rheb-GTP from PDEδ onto perinuclear membranes, combined with TSC2 mediated hydrolysis to Rheb-GDP that is resolubilized by PDEδ, generates a flux in what constitutes a spatial Rheb cycle. This spatial cycle not only counters equilibration of Rheb to all membranes, but also enables Rheb to cycle through inactivating GAP activity on TSC2-containing membranes and guanine nucleotide exchange in the cytosol. Growth factor signals that inhibit TSC2 stall the spatial cycle by, on the one hand, accumulating cytosolic Rheb-GTP on perinuclear membranes by Arl2-mediated release from PDEδ, and on the other hand, by not releasing Rheb-GTP into the cytosol due to its interaction with mTOR. Inactivation of TSC2 thereby shifts cytosolic Rheb-GTP to perinuclear membranes, resulting in mTORC1 activation. Therefore, this spatial cycle enables mTORC1 activity to be regulated solely by inhibitory signals on the GAP, TSC2 (**Fig. 7**).

**Fig. 7.**
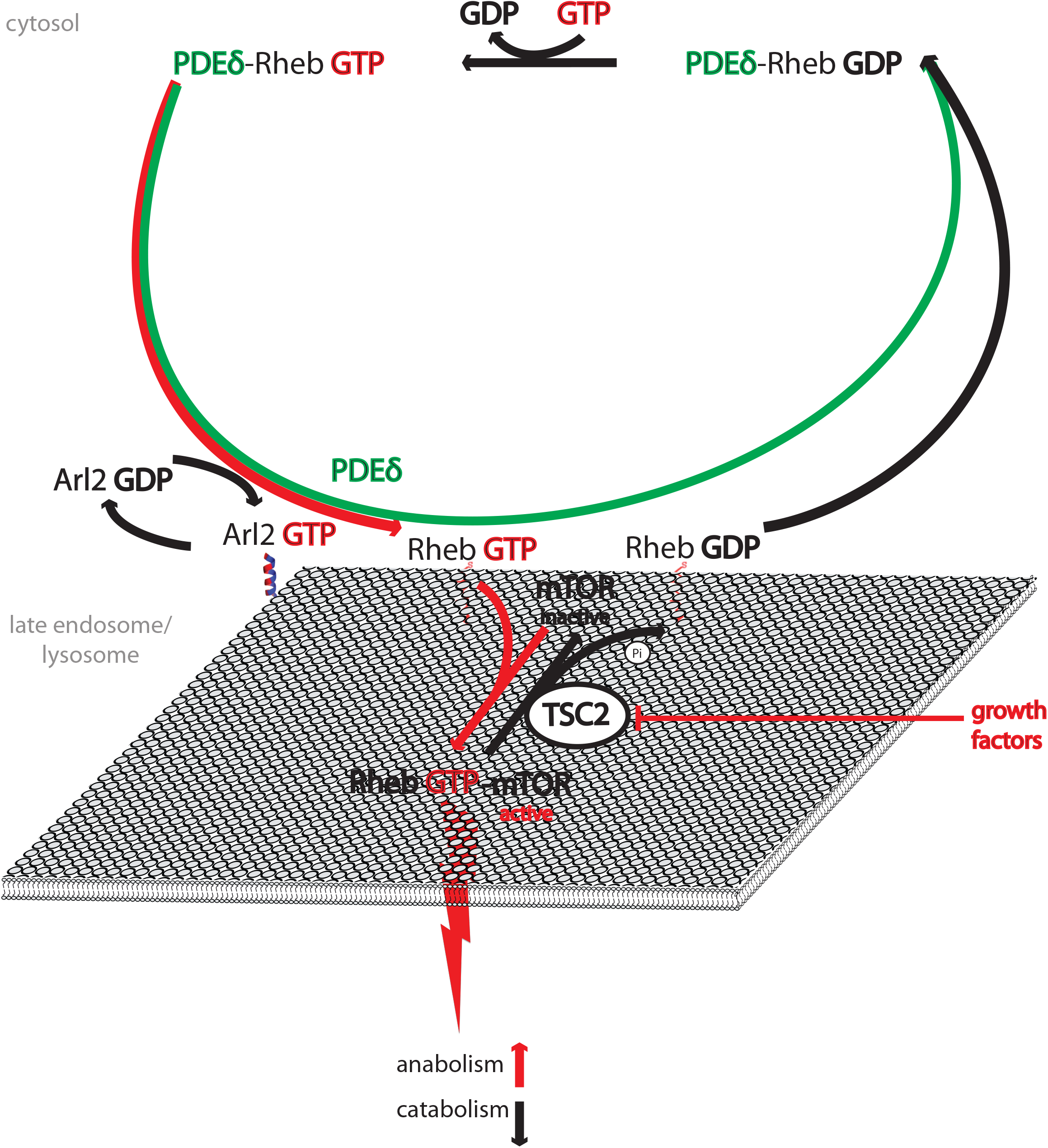
The spatially regulated GTPase cycle of Rheb. Rheb-GDP dissociating from perinuclear membranes is sequestered by PDEδ (green) in the cytosol, where efficient exchange of GDP (black) to GTP (red) occurs. The PDEδ/Rheb-GTP complex dissociates by localized Arl2-GTP activity to unload Rheb-GTP onto perinuclear membranes. On the lysosomal surface, Rheb-GTP stably binds to and activates the effector mTOR, resulting in promotion of anabolic and inhibition of catabolic processes. In the absence of growth factors, the Rheb-GAP TSC2 hydrolyzes Rheb-GTP to Rheb-GDP, which is then released from mTOR and readily dissociates from membranes to be re-sequestered by PDEδ in the cytosol.

The level of active GTP-bound Rheb in cells is higher than for most other GTPases (14, 15, 40). It was reported that in the absence of GAP activity the GTP-bound state of Rheb is maintained by an auto-inhibitory conformation with slow intrinsic GTP-hydrolysis (45). Additionally, it was reported that the intrinsic nucleotide exchange rate of Rheb is markedly higher in solution than when bound to membranes (44). A guanine nucleotide exchange factor for Rheb that could accelerate the slow nucleotide exchange rate on membranes to overcome the TSC2 GAP activity has so far not been identified. However, the data presented here suggest that the nucleotide exchange of Rheb occurs in the cytosol. The solubilizing action of PDEδ shifts Rheb-GDP from membranes to the cytosol where its intrinsic nucleotide exchange rate is higher, thereby accumulating a pool of Rheb-GTP that can rapidly activate mTOR when deposited on membranes without TSC2 activity. In the absence of growth factors, Rheb-GTP is continuously cycled through the active TSC2-containing membranes by Arl2-mediated release from PDEδ. This maintains a steady state of inactive Rheb-GDP on perinuclear membranes that cannot activate mTOR. Upon growth factor induced inactivation of TSC2, the rate limiting step of mTOR activation is now determined by the Arl2/PDEδ mediated Rheb-GTP deposition on perinuclear membranes from the cytosolic pool. Although the passive process of PDEδ binding to prenylated Ras proteins in the cytosol occurs regardless of their nucleotide-bound state (35, 46), PDEδ–mediated solubilization of Rheb, is however an indispensable factor for efficient nucleotide exchange in the cytosol.

This is further corroborated by the increased GTP/GDP ratio of Rheb in cells harboring an intact TSC2 activity and a PDEδ knock out. Without the PDEδ-mediated solubilization, Rheb is no longer enriched at perinuclear compartments by Arl2-mediated localized release, resulting in an equilibration of Rheb to all endomembranes. This withdraws Rheb from hydrolysis at TSC2 bearing perinuclear compartments. Thus, activation of the mTORC1 complex is reduced in those cells, despite an enhanced fraction of GTP-loaded Rheb.

We have shown that only Arl2, not Arl3, acts as an allosteric displacement factor of delivery of PDEδ-solubilized Rheb on perinuclear membranes, although it was reported that both Arl2 and Arl3 interact with PDEδ in a GTP-dependent manner (30). This is in agreement with previous studies that demonstrated that Arl2 knockdown was sufficient to disrupt the allosteric release of K-Ras from PDEδ (22). Structural and biochemical studies have shown that Arl3, but not Arl2 regulates the release of myristoylated ciliary proteins from different GDIs, UNC119a and UNC119b (47), further corroborating that only Arl2 displaces prenylated protein cargo from PDEδ *in vivo*. Arl2 and Arl3 interact with membranes via their N-terminal amphipathic helices (36), consistent with the observed significant enrichment of Arl2 on perinuclear membranes. Although this membrane binding is GTP-dependent for Arl3, Arl2 binds membranes in a nucleotide-independent manner (36). We have however also observed a cytosolic pool of Arl2. It was reported that cytosolic Arl2 tightly binds the tubulin-specific chaperone cofactor D, and that Arl2 in this complex is mostly GDP-bound (48). It thus is likely that Arl2 also undergoes a cytosol-membrane spatial cycle, analogous to Rheb. This leaves the question open whether an Arl2 GEF that is partitioned to perinuclear membranes regulates its activation.

We have shown that cells with an intact RTK/PI3K/TSC signaling axis (TSC2+/+ MEFs) display a controlled response to activating stimuli, such as insulin. However, insensitivity to the activating inputs and hyperactivation of mTORC1 is a hallmark of various cancer cell lines (renal, colon and ovarian cancers) (49, 50), which carry mutations targeting mTORC1 regulatory elements. Small molecules targeting mTOR in cancer, such as rapamycin, have been widely used. However, rapamycin inhibition results in the nonspecific activation of the PI3K/Akt pathway, enabling cancer cell proliferation via alternate routes (51, 52). On the other hand, disruption of PDEδ-mediated solubilization of Ras proteins, resulting in their mislocalization, was shown to be a viable strategy to inhibit proliferation of cells in Ras-driven cancer types (29, 32, 53).

PDEδ-mediated solubilization has been shown to be an essential factor in spatial cycle of Ras proteins, which maintain their subcellular localization (22, 24). Here we present a novel role of PDEδ for the Ras-related protein Rheb. Through solubilization, PDEδ not only counters equilibration of Rheb on all cell membranes, but most importantly, enables efficient nucleotide exchange on Rheb in the cytosol, leading to its reactivation.

This coupling of the generic activity of the PDEδ/Arl2-GTP mechanism to a stimulus-dependent localization of regulatory GAP enables tunable signal transduction of anabolic processes mediated by the Rheb/TSC/mTORC1 axis.

## Acknowledgements

The project was funded by the European Research Council (ERC AdG 322637) to P.I.H.B. The authors would like to thank Prof. Dr. Aurelio Teleman, Dr. Constantinos Demetriades and Prof. Dr. David Kwiatkowski for providing TSC2+/+ and TSC2-/- MEFs and Prof. Dr. Roger S. Goody for advice regarding the radioactivity experiments.

## Author Contributions

P.I.H.B., M.K and C.H.K designed the experiments. M.K. and C.H.K. performed the experiments and the analysis. M.K and A.D.K. generated the sgRNAs for PDEδ. L.R. performed western blots and the GTP/GDP assay. A.S., A.D.K. and M.K. analyzed the PLA distribution. P.I.H.B., M.K., C.H.K. and A.U.K. wrote the manuscript.

## Data availability

Data supporting the findings of this manuscript are available from the corresponding authors upon reasonable request.

## MATERIALS AND METHODS

### Materials

Primary antibodies used in this study were obtained from following sources: pS6P (S235/6) #4856 (1:1000 for WB), S6P #2317 (1:500 for WB), Rheb #13879 (1:500 for WB), TOR #2983 (1:100 for IF), Rab7 #9376 (1:100 for IF) from Cell Signaling Technology; Arl2 ab183510 (1:3000 for WB, 1:100 for IF, 1:2000 for PLA) and mCherry ab167453 (1:500 for WB) from Abcam, PDE6D H00005147-M06 (1:100 for IF, 1:1000 for PLA) and Rheb H00006009-M01 (1:100 for IF) from Abnova, Rheb NBP2-50273 (1:12.5 for IP) from Novus Biologicals, Arl3 10961-1-AP (1:300 for WB) from ProteinTech, PDE6D sc-50260 (1:300 for WB) from Santa Cruz Biotechnology, GFP #632381 (1:500 for WB) from Living Colors and α-tubulin T6074 (1:3000 for WB) from Sigma-Aldrich. Secondary antibodies for Western blot IRDye^®^680RD Donkey anti-rabbit, IRDye^®^800CW Donkey anti-mouse and IRDye^®^800CW Donkey anti-goat (1:5000 dilution) were purchased from LI-COR^®^, and secondary antibodies Alexa Fluor^®^ 647 donkey anti-rabbit A-31573 and Alexa Fluor^®^ 488 donkey anti-mouse A-21202 (1:500 dilution) used for IF were purchased from (Thermo Fisher Scientific).

Acrylamide and Precision Plus Protein^™^ standards were purchased from Bio-Rad Laboratories, Inc., APS, 2-Mercaptoethanol, NaCl, Triton X-1000 and Tween-20 from SERVA Electrophoresis GmbH, Bromophenolblue, Crystal Violet, Insulin (I9278), IGEPAL^®^ CA-630, Phosphatase Inhibitor Cocktail 2 and 3, NaDOC, TEMED and DUOLINK^®^ *In Situ* Detection Reagents FarRed (DUO92013) with PLA probes (DUO92005 and DUO92001) from Sigma Aldrich^®^, Complete Mini EDTA-free protease inhibitor tablets from Roche Applied Science, Na_2_HPO_4_ and MgCl_2_ from Merck, Ethanol, KH_2_PO_4_, KCL, Na_4_O_7_P_2_ and Tris-HCl from J.T.Baker, EDTA from Fluka^®^ Analytical, EDTA from Life Science, Glycerol, kanamycin sulfate and DTT from GerbuBiotechnik GmbH, Glycine, Roti^®^-Histofix 4%, SDS and Tris-Base from Carl Roth GmbH, Methanol from AppliChem GmbH, Micro BCA^™^ Protein Assay Kit from Thermo Scientific. ROTI^®^-Prep Plasmid mini kit for isolation of plasmid DNA was purchased from Roth, Zymoclean^™^ Gel DNA Recovery Kit from Zymo Research and NucleoSEQ^®^ Columns ansNucleoBond^®^ Xtra Midi Plus EF kit for isolation of plasmid DNA from Macherey-Nagel.

All reagents for cell culture work, DPBS, DMEM, FCS, 100x NEAA, Penicilline/Streptomycin, Trypsin/EDTA and puromycin were purchased from PAN^™^ Biotech GmbH.

Fugene^®^ 6 transfection reagent was purchased from from Promega and Lipofectamine^™^ 2000 from Invitrogen^™^ Life Technologies. Non-targeting RNA (DharmaconD-001810-10-05 ON-TARGET plus Non-targetingTargeting Pool), siRNA Arl2 (L-063126-01-0005, ON-TARGET plus Mouse Arl2 (56327) siRNA - SMART pool, 5 nmol), siRNA Arl3 (L-041895-00-0005, ON-TARGET plus Mouse Arl3 (56350) siRNA - SMART pool, 5 nmol) and DharmaFECT 1 siRNA Transfection Reagent were purchased from Dharmacon.

### Methods

#### Plasmid construction

Restriction and modification enzymes were purchased from New England Biolabs (NEB, Frankfurt am Main, Germany). PCR amplification reactions were performed with Q5 High-fidelity DNA polymerase according to the manufacturer’s instructions (NEB). All PCR-derived constructs were verified by sequencing.

mCitrine-C1 vector was generated by insertion of AgeI/BsrGI PCR fragments of mCitrine (gift from R.Tsien) cDNA into pEGFP-C1 vector (Clontech, Saint-Germain-en-Laye, France). cDNA clones encoding full-length human *RHEB* (accession number: NM005614.3) were amplified by using primers that introduce an XhoI restriction site on 5’ and EcoRI on 3’ end in mCitrine-C1 vector to generate mCitrine-Rheb. mCherry-Rheb was generated by exchanging mCitrine-C1 fluorophore to mCitrine by cutting the vector at 5’NheI and 3’ BsrGI site. Site-directed mutagenesis was performed on pEGFP-C1-Rheb to generate EYFP-Rheb Q64L.

mCitrine-Rheb HVR was generated by creating oligonucleotide corresponding to the last C-terminal 20 amino acids +CAAX-box (72 nucleotides) of full-length *RHEB*. The primers used for amplification of the PCR product were: ctcagatctcgagccaggataat (forward) and gactgcagaattctcacatcacc (reverse). The pECFP-C1 vector containing mCitrine-C1 was digested with XhoI on the 5’ end and EcoRI on the 3’ end, and ligated with Rheb-HVR amplified oligonucleotide using T4 ligase. After transformation of chemically competent XL-10 *Escherichia coli* with the ligation mix and seeding the bacterial culture on agar plate supplemented with 50 μg/mL Kanamycin, positive clones were selected for further use. All constructs were verified by sequencing. Generation of the mCitrine-Rheb and mCherry-PDEδ constructs was described previously (24).

#### Cell culture

TSC2 +/+ and TSC2 -/- MEFs were a kind gift from Prof. Aurelio Teleman (DKFZ, Heidelberg) with permission from Prof. David Kwiatkowski (Harvard). Cells were cultured in high glucose DMEM supplemented with 10% fetal calf serum, 1% non-essential amino acids, 1% L-Glutamine and 1% Pen/Strep antibiotic solution (PAN^™^ Biotech GmbH, Aidenbach, Germany) (full DMEM from here on). Cells were maintained in a humidified incubator with 5% CO_2_ at 37°C. When starvation was necessary for the experiment, cells were washed once with 1x PBS, and starved overnight in high glucose DMEM without any supplements.

#### Generation of stably transfected cell lines

The protocol for creating knockout cells via Crispr-Cas was described in (42). sgRNA was cloned into BbsI site of pSpCas9(BB)-2A-Puro (PX459) vector, Addgene number: 48139. The sgRNA sequences (5’- **CACC**TCATGTCAGCCAAGGACGAG-3’ and 3’- **AAAC**CTCGTCCTTGGCTGACATGA-5’) targeted 5’UTR region of of PDEδ mouse gene. They were generated using the online CRISPR Design Tool (http://crispr.mit.edu). The ligated sgRNA-pSpCas9 (BB)-2A-Puro plasmid, treated with PlasmidSafe was transformed in chemically competent Stbl3 bacteria on an agar plate with 100 μg/mL ampicillin. From the cultures that grew at 37°C overnight, mini- and endonuclease-free midi-preps were performed. 1*10^6^ TSC2+/+ and TSC2-/- MEFs were seeded in a 10-cm dish, and sgRNA-pSpCas9 (BB)-2A-Puro plasmid or pSpCas9 (BB)-2A-Puro (empty vector) were transfected using Fugene^®^ HD transfection reagent. The next day 2μg/mL of puromycin was applied for selection of transfected cells. The media (full DMEM supplemented with puromycin) was exchanged every 2-3 days until cells were confluent enough for subculturing and cryopreservation.

#### Transient transfection

Cells were seeded at 1*10^4^/well density in a 4-well LabTek dish or 1*10^5^/well density for 6-well dish in full DMEM. Cells were transfected with fluorescently tagged proteins at 80% confluency using Lipofectamine 2000 (Thermo Fischer Scientific, Dreiech, Germany) or Fugene^®^ HD (Promega, Mannheim, Germany) transfection reagent according to manufacturer’s guidelines.

#### Arl2 knockdown

1*10^5^ cells/well were seeded in 6-well plate or 1*10^4^/ well in a 4-well Labtek dish in DMEM supplemented with 10% fetal calf serum, 1% non-essential amino acids, 1% L-Glutamine and 1% Pen/Strep antibiotic solution. Cells were transfected at 80% confluency with 50nM of non-targeting RNA, siRNA Arl2 or Arl3 by use of DharmaFECT 1 siRNA Transfection Reagent according to manufacturer’s protocol (GE Dharmacon, Lafayette, USA). 7-10 hours post-transfection the medium was changed to full DMEM and cells were transfected with 0.25 μg of mCitrine-Rheb cDNA. The image collection and cell-lysis was performed 48 hours post siRNA transfection.

#### Microscopy

Microscopic Imaging was carried out on a confocal microscope Leica TCS SP5 (Leica Microsystems), Olympus FluoroView FV1000 (Olympus) or wide field CellR (Olympus). Images from the Leica TCS SP5 were obtained at 512*512 pixels at 400 Hz, with a 63*1.4 N.A. objective. The pinhole size was set to 250 μm and fluorescence was detected with a photomultiplier tube (PMT) set at 1225 V. The 514 nM argon laser line at 30% power was used to excite mCitrine-Rheb and fluorescence was collected in the 525-625 nm spectral range. Images were acquired with 3 times line averaging and obtained every 2 minutes for 1 hour.

On the Olympus FluoroView FV1000, DAPI/Hoechst was excited with 405 nm, mTFP with 458 nm, mCitrine and Alexa488 with the 488 nm argon laser lines. mCherry was excited with the 561 nm line of a DPSS laser, and Alexa 647 with the 633 nm line of a HeNe laser. PLA puncta were imaged through a 40x /0.9 N.A. air objective, whereas a 60x /1.2 N.A. oil immersion objective was used otherwise. Fluorescence excitation/emission was selected through the dichroic mirrors DM 458/515 and DM 405/488/561/633, with the pinhole set at 250 μm. Fluorescence spectral selection was through an acousto-optic tunable filter (AOTF) and SIM scanner. A 420-460 nm bandwidth was set for DAPI fluorescence detection, 498-552 nm for Alexa488 and mCitrine, 571-650 nM for mCherry and 655-755 nm for Alexa 647. Sequential imaging was performed with 3 averaged frames and by application of a Kalman filter.

Wide field images acquired on the CellR (Olympus) were obtained at 672*512 pixels with a 60x 1.2 N.A. oil immersion objective. DAPI fluorescence was obtained through a TagBFP+ filter and far-red fluorescence through a cy5 filter at 66ms exposure time.

In all cases, live cell imaging was performed at 37°C with 5% CO_2_, whereas fixed samples were imaged at room temperature.

#### Image processing

Average background fluorescence was obtained from a cell free region and subtracted from all images. For time lapse series, a bleach correction was performed by normalizing the total image intensity of a frame to the average intensity of the starting image. Quantification of co-localization of Rab7/mTOR with endogenous Rheb or mCitrine-Rheb was done by masking the fluorescence of Rab7/mTOR by intensity-based thresholding. The ratio of Rheb/mCitrine-Rheb fluorescence in the resulting region of interest over total Rheb/mCitrine-Rheb fluorescence in the cells was then used as measure of the fraction of Rheb that localizes to Rab7/mTOR positive membranes.

A relative estimate of the soluble fraction of mCitrine-Rheb was obtained by calculating the nuclear over total mCitrine-Rheb fluorescence intensity. The perinuclear fraction was obtained by performing intensity-based thresholding on cells expressing mCitrine-Rheb and calculating the ratio of perinuclear over total cell mCitrine-Rheb intensity.

#### Fluorescence Lifetime Imaging Microscopy (FLIM)

Fluorescence lifetime images were acquired using a confocal laser-scanning microscope (FV1000, Olympus) equipped with a time-correlated single-photon counting module (LSM Upgrade Kit, Picoquant). For detection of the donor (mCitrine), the sample was excited using a 510 nm diode laser (LDH 507, Picoquant) at 36 MHz repetition frequency. Fluorescence signal was collected through an oil immersion objective (60x/1.35 UPlanSApo, Olympus) and spectrally filtered using a narrow-band emission filter (HQ 530/11, Chroma). Photons were detected using a single-photon counting avalanche photodiode (PDM Series, MPD) and timed using a single-photon counting module (PicoHarp 300, Picoquant). Data were analyzed by global analysis (34).

#### Immunofluorescence

One day post-transfection or seeding, cells were washed once with PBS, fixed with Roti^®^-Histofix (Carl Roth GmbH, Karlsruhe, Germany) for 10 minutes at RT and permeabilized for 10 minutes with PBS+0.1% Triton X-100 (SERVA Electrophoresis GmbH, Heidelberg, Germany). The blocking was performed by applying Odyssey^®^ Blocking Buffer (LI-COR Biosciences GmbH, Bad Homburg vor der Höhe, Germany) for 1h at RT. Primary antibodies were diluted at the recommended ratio (see Materials section) in Odyssey^®^ Blocking Buffer and incubated overnight at 4°C. The next day, following 3×5minutes washing with PBS+0.1% Tween-20 (SERVA Electrophoresis GmbH, Heidelberg, Germany), secondary antibodies were prepared in Odyssey^®^ Blocking Buffer and incubated for 1h at RT. After final washing steps with PBS+ 0.1% Tween-20 of 3×5 minutes, PBS was added to cells and the cells were stored at 4°C until imaging.

#### Proximity Ligation Assay (PLA)

8000 cells/well were seeded in an 8-well Labtek in full DMEM. The next day, the cells were once washed with PBS and fixed with Roti^®^-Histofix for 10 minutes, permeabilized with PBS+0.1% Triton X-100 for 10 minutes and blocked with a blocking solution (DUO92005-100RXN, Sigma-Aldrich^®^, Taufkirchen, Germany) for 1h at RT. Arl2 and PDEδ antibodies were diluted in Antibody diluent (DUO92005-100RXN, Sigma-Aldrich^®^, Taufkirchen, Germany), and applied separately (negative control) and together in individual wells, and incubated overnight at 4°C. The next day, the Duolink *in situ* PLA assay (DUO92013-100RXN, Sigma-Aldrich^®^, Taufkirchen, Germany) was performed according to manufacturer’s instructions. Finally, the nuclei were stained with DAPI (Sigma-Aldrich^®^, Taufkirchen, Germany) diluted 1:500 in PBS for 15 minutes. The cells were stored at 4°C in PBS until imaging.

#### PLA distribution analysis

For the quantification of PLA puncta distributions, we calculated the distance between each punctum in 3-D confocal image stacks and the respective nuclear center. PLA puncta were identified using the Trackpy module (github.com/soft-matter/trackpy, DOI 10.5281/zenodo.60550) in the Anaconda Python programming language (Python Software Foundation, version 2.7, https://www.python.org/). The nuclear envelope was calculated to estimate the nuclear center, mean nuclear radius as well as to identify which PLA puncta are localized in the nucleus. Analogous analysis was performed on the 2-D projections on the focal plane for more accessible comparison to a reference random distribution, which was generated by calculating the distance of each pixel from the 2-D cell mask to the nuclear center. This embodies in essence a Monte Carlo approach, where each of the PLA puncta is randomly redistributed throughout the pixels of the cells mask multiple times with the final distribution being the estimated average distribution to the nuclear radius. Consequently, the cell shape and size determine the random distance distribution and therefore have influence on the sensitivity of the identification of perinuclear localization. Cells with less than 10 PLA puncta were excluded from the analysis.

Spatial bins represented in graphs in **Supplementary Fig. 1b** were created by subtracting values of the randomized puncta distribution from the experimentally obtained one. The size of individual bins corresponds to the division of the maximal size of the cell with the number of wanted bins.

#### Segment analysis

The analysis was made with an in-house developed software developed in Anaconda Python (Python Software Foundation, version 2.7, https://www.python.org/).

For the analysis of fluorescence intensity distributions in cells, nuclear and total cell masks for every cell were created in ImageJ 1.47k (https://imagej.nih.gov/ii/)

For each pixel in the cell, the distance *(d)* to the nuclear membrane was normalized:

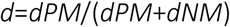

Where *dNM* is the shortest distance to the nuclear membrane and *dPM* the shortest distance to the plasma membrane.

According to the calculated distances, the cells were divided in 3 spatial bins (1 being closest to the plasma membrane and 3 being closest to the nuclear membrane), with equal radius between them (segment thickness identical across the cell). For angular masking, the center of the nucleus was determined and the intensity weighted ‘center of the cell’ and a central axis was fitted to these two points. An 60° angle around this axis was used for the analysis of the intensity distributions. Images for segmentation were defined to appear in a designated order, with a corresponding mean intensity profile for each segment. Each value of the segment was normalized to the sum of intensity in all segments, and the resulting mean value with the standard deviation was plotted in the graph. P values were determined by a Student’s unpaired t-test. All values <0.05 were determined as not significant.

#### Western blots

TSC2+/+ MEFs were seeded in 1*10^5^, and TSC2-/- MEFs in 8*10^4^ density in full DMEM in a 6-well plate. The next day the cells were starved overnight and the following day treated with 300 nM insulin (Sigma-Aldrich^®^, Taufkirchen, Germany) for 15 minutes before preparation of whole cell lysates, unless indicated differently. Cells were lysed with ice-cold RIPA buffer supplemented with Complete Mini EDTA free protease inhibitor (Roche Applied Science, Penzberg, Germany) and phosphatase inhibitor cocktail 2 and 3 (Sigma-Aldrich^®^, Taufkirchen, Germany), scraped after 10 minutes of incubation on ice and passed through a 26G needle 5-7 times followed by centrifugation at 14 000 rpm, 4°C for 15 minutes. Protein concentration was determined using BCA assay, with BSA in different concentrations used as the standard. 25μg of whole cell-lysate was used for the SDS-PAGE. The gels were blotted to a polyvinylidene difluoride (PVDF) membrane, which was pre-activated with methanol for 5 minutes. After successful transfer, blocking was performed for 1 h by placing the membrane in Odyssey Infrared Imaging System blocking buffer. The antibodies were diluted in blocking buffer and incubated overnight at 4°C (for antibodies used and dilutions see section Materials). The next day, membranes were washed 3×5 minutes with 1xTBST (+ 0.1% Tween-20), followed incubation with secondary antibodies (diluted in blocking buffer) for 1h at RT. The membranes were again washed 3×5 minutes with 1x TBST (0.1% Tween) and protein detection was done with Odyssey Imaging System.

#### Real-Time Cell Analyzer (RTCA)

RTCA was performed using 16-well E-plates on the Dual Plate xCELLigence instrument (Roche Applied Science, Indianapolis, IN, USA). This system measures a dimensionless parameter called cell index (CI), which evaluates the ionic environment at an electrode/solution interface and integrates information on cell number. 5000 cells of TSC2 +/+, TSC2 +/+ E.V. and TSC2 +/+ sg RNA PDEδ MEFs and 3500 of TSC2 -/-, TSC2 -/- E.V. and TSC2p53-/- sg RNA PDEδ MEFs were plated in each well of the 16-well plates in 200 μL of full DMEM. After seeding, cells were allowed to settle for 30 min at room temperature (RT) before being inserted into the xCELLigence instrument in a humidified incubator at 37°C with 5% CO_2_. Impedance measurements were then monitored every 15 min up to 200 hours. All assays were performed in duplicates. The growth rate was analyzed by calculating the slope of the curve between 40 and 60 hours by using the following formula:

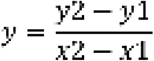

where y represents the average growth rate (slope) with y2 the measured cell density at 60-hours (x2) and y1 the measured cell density at 40 hours (x1).

#### Colony formation assays

100 cells/well were seeded in 6-well plate in full DMEM and incubated in a humidified incubator with 5% CO_2_ at 37°C for 10 days. The medium was changed every 2 to 3 days to avoid deprivation of nutrients. Finally, cells were once washed with PBS, fixed with Roti^®^-Histofix for 10 minutes at RT, washed 3x with PBS and incubated with 0.01% (v/v) Crystal Violet (Sigma-Aldrich^®^, Taufkirchen, Germany). After 1 hour, Crystal Violet was aspirated; the wells were washed 2-3 times with ddH_2_O and left for drying. The plates were scanned with Typhoon TRIO + scanner (Amersham Biosciences, Little Chalfont, UK) and analyzed using a script from (54).

#### Nucleotide loading state of Rheb

Radioactive nucleotide labeling was performed in a variation of the method described by (55). In brief, cells were starved for 18h in phosphate-free DMEM. Then, cells were washed and incubated for 5h in phosphate-free DMEM containing 0.15 mCi [32P]-orthophosphate. Afterwards, cells were lysed with lysis buffer (50 mM Tris pH 7.4, 140 nM NaCl, 1% Triton X-100, 1% IGEPAL CA-360, 1 mM KCl, 2 mM MgCl_2_, 1x EDTA-free protease inhibitor cocktail) and centrifuged for 15 min, 4°C, 13,000 rpm. Lysates were pre-cleared with Protein A-Sepharose before immunoprecipitation of Rheb with an anti-Rheb antibody bound to Protein A-Sepharose beads (ratio 1:12.5) for 45 min at 4°C on a rotator. Immunoprecipitates were washed six times with washing buffer (50 mM Hepes pH 7.4, 500 nM NaCl, 0.1% Triton X-100, 0.005% SDS, 5 mM MgCl_2_). Afterwards, supernatant was removed with a syringe (G27) and GTP/GDP were eluted from the beads with elution buffer (2 mM EDTA, 0.2% SDS, 1 mM GDP, 1 mM GTP) for 20 min at 68°C. To separate GTP and GDP fractions, thin layer chromatography (TLC) was performed on PEI-cellulose plates developed in 1 M KH_2_PO_4_ pH 3.4. Diluted [alpha-32P]-GTP, either untreated or treated with 0.001 units of Shrimp Alkaline Phosphatase (NEB) on ice, were spotted as reference on every plate. Afterwards, dried plates were exposed to storage phosphor screens for 5 − 8 days and scanned on a Typhoon Trio Imager. The intensity profile of each TLC lane was plotted using FiJi and analyzed by Multi-peak fitting and background correction using IGOR Pro, version 6.37. The resulting intensity values for the GTP and GDP spots were divided by the factor three and two, respectively, to account for the number of phosphate groups per nucleotide, and the percentage of GTP was determined as GTP/(GTP+GDP)*100%.

## Supplementary files

Supplementary information contains Supplementary Fig. 1-9

